# Wnt signaling activates gene expression in the absence of the *C. elegans* DREAM repressor complex in somatic cells

**DOI:** 10.1101/811414

**Authors:** Jerrin R. Cherian, Lisa N. Petrella

## Abstract

Establishment and maintenance of proper gene expression is a requirement for normal growth and development. The DREAM complex in *Caenorhabditis elegans* functions as a transcriptional repressor of germline genes in somatic cells. At 26°C, DREAM complex mutants show temperature associated increase in misexpression of germline genes in somatic cells and High Temperature Arrest (HTA) of worms at the first larval stage. To identify transcription factors required for the ectopic expression of germline genes in DREAM complex mutants, we conducted an RNA interference screen against 123 transcription factors capable of binding DREAM target promoter loci for suppression of the HTA phenotype in *lin-54* mutants. We found 15 embryonically expressed transcription factors that suppress the HTA phenotype in *lin-54* mutants. Five of the transcription factors found in the initial screen interact with the Wnt signaling pathways. In a subsequent RNAi suppression screen of Wnt signaling factors we found that knock-down of the non-canonical Wnt/PCP pathway factors *vang-1*, *prkl-1* and *fmi-1* in *lin-54* mutant background resulted in strong suppression of the HTA phenotype. Animals mutant for both *lin-54* and *vang-1* showed almost complete suppression of the HTA phenotype, *pgl-1* misexpression, and fertility defects associated with *lin-54* single mutants at 26°C. We propose a model whereby a set of embryonically expressed transcription factors, and the Wnt/PCP pathway, act opportunistically to activate DREAM complex target genes in somatic cells of DREAM complex mutants at 26°C.

## INTRODUCTION

Properly regulated gene expression is critical for normal growth and development of an organism. In metazoan development, embryonic cells become either totipotent germ cells, or somatic cells that undergo cell fate specification and differentiation. Consequently, somatic cells must repress the gene expression program associated with germline cells. Mis-regulation of gene expression control is associated with altered cell-fate specification, which can lead to retarded development or diseases such as cancer (Gross *et al*. 2016; Smith *et al*. 2019). For example, hundreds of genes expressed primarily in germ cells (germline genes) have been found to be up-regulated in 19 different somatic cancers (Wang *et al*. 2016). Evidence suggests that germline genes can potentially function to drive cancer acquisition and progression (Chang *et al*. 2019). To maintain correct cell fate, precise control of both spatial and temporal gene expression is required via a network of transcriptional activators and repressors (Kudron *et al*. 2013).

In *C. elegans*, the DREAM complex alters chromatin structure in the nucleus to transcriptionally repress germline genes in somatic cells (Costello and Petrella 2019; Cui *et al*. 2006; Latorre *et al*. 2015; Petrella *et al*. 2011; Rechtsteiner *et al*. 2019; Unhavaithaya *et al*. 2002; Wang *et al*. 2005; Wu *et al*. 2012). The DREAM complex is completely conserved between mammals and *C. elegans*, and comprised of eight proteins: E2F-DP heterodimer (EFL-1 & DPL-1), a retinoblastoma-like pocket protein (LIN-35), and a 5-subunit MuvB complex (LIN-9, LIN-37, LIN-52, LIN-53, and the DNA binding subunit LIN-54) (Goetsch *et al*. 2017; Harrison *et al*. 2006; Latorre *et al*. 2015; Sadasivam and DeCaprio 2013). In *C. elegans*, a mutant form of LIN-54 is incapable of binding the majority of DREAM complex target loci due to a change in a single amino acid in the DNA binding motif (Tabuchi *et al*. 2011). A putative null allele of *lin-35*, which physically mediates the interaction between the E2F-DP dimer and muvB subcomplex, results in reduced DREAM complex binding to target loci and reduced repression capability of the DREAM complex (Goestsch *et al*. 2017). Mutation in *lin-54*, *lin-35*, and several other members of the DREAM complex members results in low level misexpression of germline genes, such as P-granules, in somatic cells at 20°C (Wang *et al*. 2005) and increased misexpression of germline genes under moderate temperature stress of 26°C (Petrella *et al*. 2011). Ectopic P-granule expression in the intestine of DREAM complex mutants is correlated with the High Temperature Larval Arrest (HTA) phenotype, where worms grown at 26°C developmentally arrest at the L1 larval stage (Petrella *et al*. 2011).

Wild-type embryonic development is characterized by rapid chromatin compaction, with cells achieving closed chromatin structures between the 100 to 200 cell stage (Mutlu *et al*. 2018). While wild type embryos show a moderate delay in this process at high temperature; DREAM complex mutants at high temperature show severe delays in the chromatin compaction process. This delay results in open chromatin persisting later in development, most prominently in an anterior embryonic cell (Costello and Petrella 2019). Open chromatin structures have been shown to facilitate recruitment of DNA binding proteins such as transcription factors (Heinz *et al*. 2011). Transcription factors act both spatially and temporally within the chromatin landscape to modulate gene expression (Li *et al*. 2018; Praggastis *et al*. 2017). Repressor complexes, such as the DREAM complex, work antagonistically to gene activation by preventing RNA polymerase access to target promoter loci (Hernández-Arriaga *et al*. 2009). The delayed chromatin compaction seen in DREAM complex mutants may provide an opportunity for transcription factors to ectopically activate germline genes in the soma. Additionally, the A-P pattern of decreased chromatin compaction in DREAM complex mutants suggests that there may be a role for cell-signaling pathways involved in embryonic A-P patterning in ectopic germline gene expression. Neither the transcription factors nor cell signaling pathways necessary for ectopic germline gene expression in DREAM complex mutants are known.

In this study, we conducted an RNAi screen of 123 transcription factors (TFs) that were predicted to bind DREAM target loci and scored for suppression of HTA in *lin-54* mutants. We found that knock-down of 15 of the transcription factors resulted in suppression of the HTA phenotype in *lin-54* mutants. Knockdown of nine of 15 TF HTA suppressors was also able to suppress the ectopic expression of the germline gene PGL-1 in *lin-54* mutants. Signaling pathway enrichment analysis of the HTA transcription factor suppressors showed significant enrichment of the Wnt signaling pathway, the primary A-P patterning signaling pathway in *C. elegans*. We therefore performed RNAi of Wnt pathway components for their ability to suppress the HTA phenotype in *lin-54* mutants. While loss of a number of Wnt components suppressed the HTA phenotype, knock-down of Wnt/PCP pathway members, including *vang-1*, showed strong suppression of HTA. *lin-54; vang-1* double mutants showed complete suppression of HTA phenotype and reduced ectopic PGL-1 expression to baseline levels. Loss of *vang-1* also completely suppressed the fertility defects seen in *lin-54* mutants. We propose that the Wnt/PCP pathway can act to promote gene expression in both the soma and germline in the absence of DREAM complex.

## MATERIALS AND METHODS

### Strains and nematode culture

*C. elegans* were cultured under standard conditions (Brenner 1974) on NGM plates seeded with *E. coli* strain AMA1004 at 20°C unless otherwise noted. N2 (Bristol) was used as wild-type. Mutant strains used for the experiments were MT8841 *lin-54(n2231),* MT10430 *lin35(n745),* MT8838 *lin-13(n770),* RB1125 *vang-1(ok1142)*, LNP0073 *lin-54(n2231);vang-1(ok1142)*, CF1045 *muIs49* [*unc-22(+), Pegl-20::egl-20::GFP*], and LNP0096 *lin-54(n2231);muIs49* [*unc-22(+), Pegl-20::egl-20::GFP*]. All strains, except LNP0073 and LNP0096 which were made for this study, were provided by the CGC, which is funded by NIH Office of Research Infrastructure Programs (P40 OD010440).

### Selection of Transcription Factor Candidates for RNAi Screen

The Yeast-1-Hybrid (Y1H) Assay dataset (Reece-Hoyes *et al*. 2011) was used to obtain a list of transcription factors (TFs) to screen for suppression of HTA phenotype in *lin-54* mutants. This dataset includes the interactions of 296 predicted transcription factors (Bass *et al*. 2016) as prey screened to interact with 534 promoter loci bait sequences (Reece-Hoyes *et al*. 2011). We found that 48 bait sequences in the Y1H dataset were known to be bound by the DREAM complex in their promoter region in late embryos (Goetsch *et al*. 2017). We found that 123 of the 296 TFs showed binding to at least one of the 48 DREAM target promoter loci in the Y1H data set. These 123 TFs were used as candidates in the transcription factor RNAi screen.

### Suppression of High Temperature Larval Arrest (HTA) Phenotype assay

Hermaphrodites were maintained at 20°C on NGM plates with AMA1004 bacteria until the L4 larval stage. At the L4 larval stage P0 worms were moved to plates containing transcription factor (TF) specific RNAi feeding bacteria NGM (0.2% lactose, 0.075 mg/ml ampicillin), and simultaneously upshifted to 26°C. After ∼ 18 hours the P0 worms were moved to new RNAi plates and allowed to lay embryos for 24 hours. For preliminary qualitative HTA suppression RNAi screen, each RNAi plate was scored as either primarily having F1 worms that were arrested at the L1 stage or primarily having worms that grew beyond L1 stage. For quantitative HTA suppression RNAi screen against TF and Wnt signaling components, individual F1 worms were counted either as arrested at L1 stage or growing to the L4/Adult stage. Worms grown on empty L4440 vector containing bacteria was used as a RNAi negative control. RNAi against *mrg-1*, a chromatin modifier previously shown to suppress HTA in DREAM mutants (Petrella *et al*. 2011), was used as a positive control. For suppression of HTA phenotype assay in *lin-54; vang-1* mutants, the assay was performed as described above except that worms were grown on regular NGM plates seeded with AMA1004 throughout the assay. Statistical significance was determined by comparing test samples with negative control using Fishers Exact test in Graph Pad Prism.

### Immunostaining

L4 stage worms were upshifted to 26°C on RNAi feeding bacteria containing plates as described above. On the next day, gravid adult worms from these plates were placed in 100µL M9 buffer on a slide in a humid chamber at 26°C to lay embryos. L1 worms were placed on a poly-L-lysine-coated slide and a coverslip was placed over the sample. Excess M9 buffer was wicked away and the slide was placed in liquid nitrogen for a minimum of five minutes. Slides were freeze cracked and fixed in cold methanol for 10 minutes followed by cold acetone for 10 minutes each. Slides were air dried and blocked for an hour with 1.5% of Bovine Serum Albumin and 1.5% of Ovalbumin in PBS (adapted from Strome and Wood 1983). Polyclonal rabbit anti-PGL-1 antibody (Kawasaki *et al*. 1998) was used at 1:10,000 dilution overnight at 4°C. Slides were washed three times in PBS for 10 minutes and blocked for 15 minutes at room temperature. Slides were incubated with secondary antibody conjugated to Alexa Fluor 647 (Thermofisher Scientific, Cat No. A-21244) at 1:500 dilution for 2 hours at room temperature. Slides were treated with 5mg/ml DAPI in 50 mL PBS for 10 minutes, washed 3 times with PBS for 10 minutes at room temperature and mounted on gelutol mounting medium. Z-stacks were taken using Nikon A1R Inverted Microscope Eclipse T*i* confocal microscope with NIS Elements AR 3.22.09 at 60X. PGL-1 quantification was done in NIS-Elements Analysis program. First, all somatic cells were outlined to separate them from primordial germ cells and create a region of interest. Second, the mean somatic PGL-1 intensity for each worm was obtained using the ROI statistics function. The adjusted PGL-1 intensity was calculated by subtracting a worm’s mean PGL-1 pixel intensity from the mean background pixel intensity and then dividing the result by mean background pixel intensity (Zhao *et al*. 2018). Statistically significant difference between test samples and L4440 empty vector negative control was calculated using Kolmogorov-Smirnov test using Graph Pad Prism. In the case of *lin-54; vang-1* mutants, worms were grown on regular NGM plates seeded with AMA1004 and the staining and analysis were performed as described above.

### Gene Ontology Enrichment Analysis

Gene Ontology Analysis was performed using two different enrichment analysis tools: gProfiler - https://biit.cs.ut.ee/gprofiler/gost (Version: e96_eg43_p13_563554d) (Raudvere *et al*. 2019) and PANTHER Overrepresentation Test - http://geneontology.org/ (Release: 20190711) (Ashburner *et al*. 2000; Mi *et al*. 2019). Transcription factor candidates that when knock-downed in *lin-54* mutants show a significantly different level of HTA suppression from negative control were uploaded to both the enrichment analysis tools and the significantly enriched ontology terms were obtained (Multiple testing correction: PANTHER - FDR<0.05; gProfiler - Custom g:SCS Set Counts and Sizes correction method, Cut off value:0.05).

### Brood Size Assay

The brood size of worms was determined at 3 different temperature conditions, 20°C, up-shift from 20°C to 26°C at the L4 stage, and 26°C. For 20°C, individual P0 hermaphrodites were placed on a plate starting at the L4 stage and moved to a new plate every 24 hours until the worms stopped having progeny. The total number of F1 progeny for each P0 worm was counted. For upshifting, P0 hermaphrodites were maintained at 20°C until the L4 stage and then shifted to 26°C at the L4 stage on individual plates. The worms were moved to a different plate every 24 hours until the worms stopped having progeny. The total number of F1 progeny for each P0 hermaphrodite were counted. For 26°C, P0 hermaphrodites were maintained at 20°C until the L4 stage and then shifted to 26°C and allowed to have progeny. The F1 progeny that had grown continuously at 26°C were placed on an individual plate at the L4 stage and moved to a new plate every 24 hours until the worms stopped having progeny. The total number of F2 progeny for each F1 worm was counted. Brood sizes of zero were excluded. Statistical analysis of brood size was done using Welch’s T test on fertile animals using Graph Pad Prism.

### EGL-20::GFP Expression Analysis

L4 stage worms were used at 20°C or upshifted to 26°C on NGM plates containing AMA1004 bacteria. On the next day, gravid adult worms from these plates were placed in 100µL M9 buffer on a slide in a humid chamber at either 20°C or 26°C to lay embryos. L1 worms were moved to a poly-L-lysine-coated slide and a coverslip was placed over the sample. M9 buffer present in excess was wicked away and the slide was kept in liquid nitrogen for a minimum of five minutes. Slides were freeze cracked, and fixed in cold methanol for 10 minutes followed by cold acetone for 10 minutes each (adapted from Strome and Wood,1983). Slides were treated with 5mg/ml DAPI in 50 mL PBS for 10 minutes, washed 3 times with PBS for 10 minutes at room temperature and mounted on gelutol mounting medium. Z-stacks were taken using Nikon A1R Inverted Microscope Eclipse T*i* confocal microscope with NIS Elements AR 3.22.09 at 60X. ROI was manually determined by picking a point on dorsal posterior portion of the worm with EGL-20:GFP intensity above a cut off value and another point farthest away the dorsal posterior portion with a cut off same value. We drew a polygonal line using measure length function in annotation and measurement function of NIS Elements Analysis program. The same analysis was performed on the ventral posterior portion of the worm. Statistical analysis was done using Students T test using Graph Pad Prism.

## RESULTS

### Multiple transcription factors are required for the high temperature larval arrest phenotype of lin-54 mutants

Loss of the DREAM complex results in up-regulation of many genes including ectopic somatic expression of genes normally only expressed in the germline (Petrella *et al*. 2011, Wang *et al*. 2005, Wu *et al*. 2012). The up-regulation of DREAM target genes in DREAM complex mutants correlates with the High Temperature larval Arrest (HTA) phenotype, characterized by a developmental arrest at the L1 stage when animals are grown at 26°C. We predicted that ectopic expression of DREAM target genes in DREAM mutants likely requires the activity of transcription factors (TFs) (Figure 1a). To find the TFs necessary for driving DREAM target gene misexpression, we performed an RNAi screen of TFs in *lin-54* mutants for suppression of the HTA phenotype. For our screen, we used a hypomorphic *lin-54*(n2231) allele as it limits DREAM complex DNA binding but is healthy and fertile. To limit the screen to TFs that directly bind to and regulate DREAM complex targets, we developed a candidate set of 123 TFs from those shown to bind DREAM targets *in vitro* by Yeast-1-Hybrid (Reece-Hoyes *et al*., 2011). For the initial screen, we performed RNAi against 123 TFs in *lin-54* mutants at 26°C and screened for the suppression of the HTA phenotype using a binary (yes/no) method where plates were scored for the presence of L4/adults worms (Figure 1b, Materials and Methods). As a positive control, we performed RNAi against *mrg-*1, a chromatin associated factor previously shown to suppress the HTA phenotype (Petrella *et al*. 2011). We found that RNAi against 26 of 123 TF candidates tested were able to suppress the HTA phenotype in *lin-54* mutants. We retested the 26 TFs by quantifying the percentage of worms that grew to the L4/adult stage. RNAi against 15 of 26 TFs (henceforth called TF HTA suppressors) showed statistically significant suppression of HTA in *lin-54* mutants compared to the empty vector L4440 RNAi control (Figure 1c, Table 1, Figure S1). The TF HTA suppressors fell into three transcription factor DNA binding domain families: homeodomain (HD), Winged Helix (WH), and zinc finger (ZF) (Table 1).

**Figure 1:**
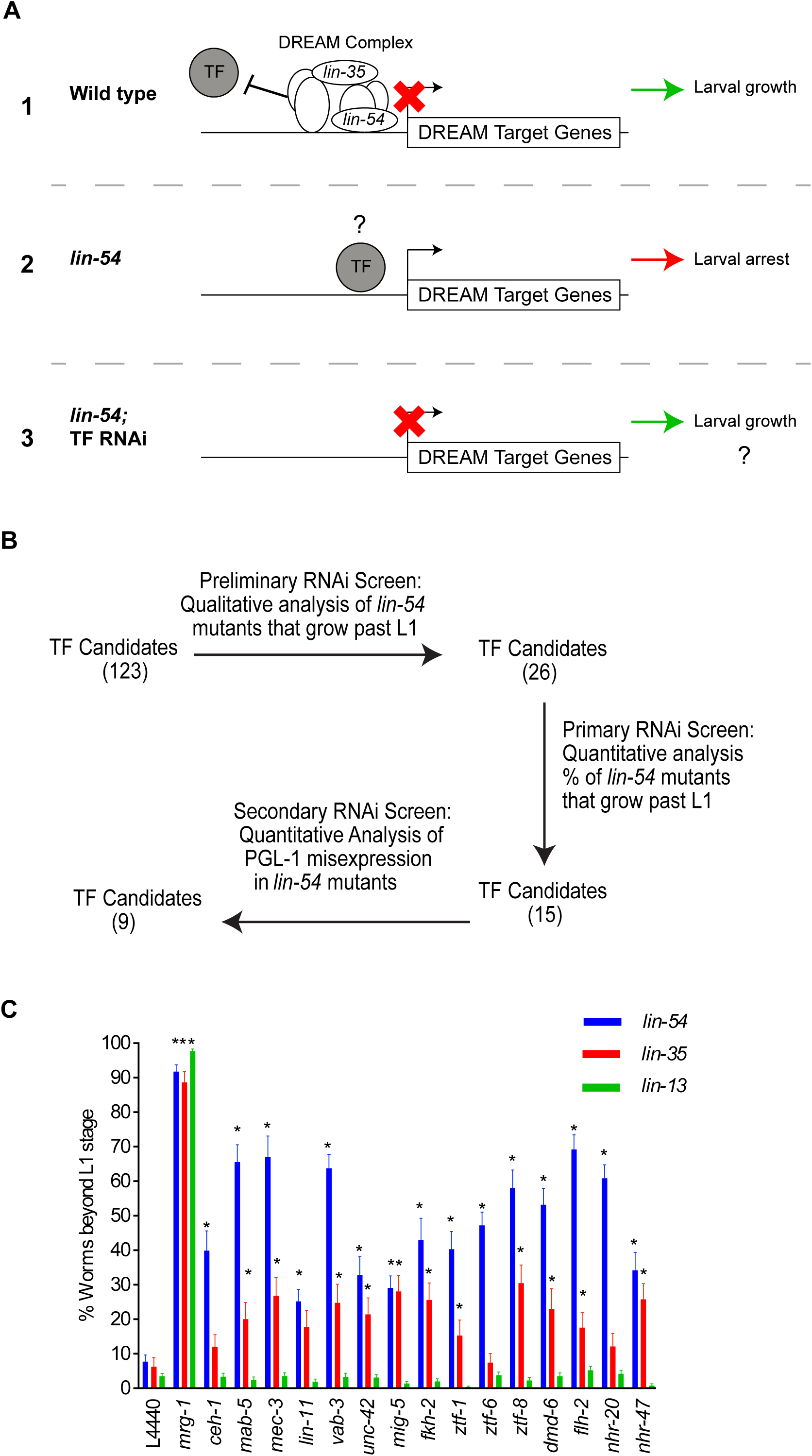
RNAi screen identifies 15 transcription factors that promote the HTA phenotype in DREAM complex mutants. (A) Model of DREAM complex target gene regulation (1) In the wild type, the DREAM complex represses germline gene misexpression in somatic cells and allows normal worm growth past the L1 stage at 26°C. (2) In *lin-54* mutants the DREAM complex cannot bind, which results in activation of DREAM target genes and the high temperature larval arrest (HTA) phenotype is observed at 26°C. We predict that under these conditions transcription factors bind (TF) to and activate DREAM target genes. (3) Therefore, when those TFs are knock-down by RNAi in *lin-54* mutant background there should be a decrease in DREAM target gene expression and restored larval growth. (B) Workflow for RNAi screen to identify transcription factors involved in suppression of HTA phenotype and reduction in germline PGL-1 misexpression in somatic cells of *lin-54* mutants. (C) Knock-down of 15 and 11 transcription factors in *lin-54* and *lin-35* mutants respectively resulted in significant suppression of the HTA phenotype when compared to L4440 empty vector RNAi. There was no suppression of the HTA phenotype in *lin-13* mutants upon knock-down of any of the 15 transcription factors. RNAi against *mrg-1* was used as a positive control for HTA suppression. Significant percentage of growth to the L4/Adult stage compared to L4440 empty vector was determine by Fisher’s Exact test (\**P* value ≤ 0.05). Error bars indicate standard error of proportion.

**Table 1:**
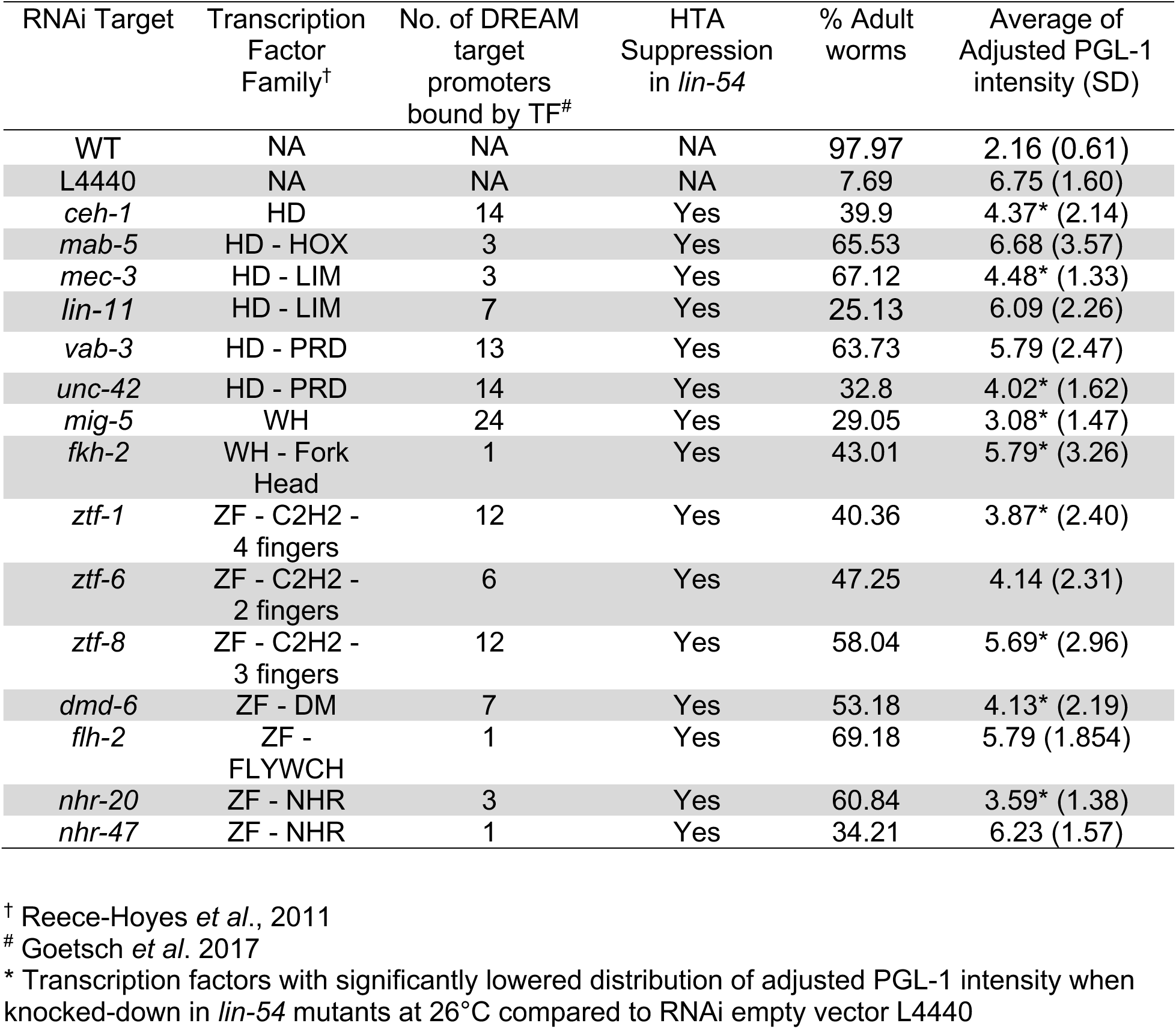
Transcription factors that suppress HTA and PGL-1 ectopic expression.

### TF HTA suppressors are required for the HTA phenotype of lin-35 mutants but not lin-13 mutants

Loss of *lin-35*, the sole Retinoblastoma homolog DREAM complex member in *C. elegans*, results in ectopic germline gene expression and the HTA phenotype (Petrella *et al*. 2011; Wang *et al*. 2005). We performed RNAi against the 15 TF HTA suppressors in a *lin-35* putative null mutant background to determine if the TFs found to suppress the HTA phenotype of *lin-54* mutants can do the same in *lin-35* mutants. RNAi against 11 of the 15 TF HTA suppressors genes in *lin-35* mutants showed significant suppression of HTA phenotype compared to empty vector RNAi (Figure 1c). Suppression of the HTA phenotype was generally weaker in *lin-35* mutants compared to *lin-54* mutants. Because lin-35 mutants have been shown to demonstrate enhanced somatic RNAi, we do not think these weaker phenotypes are due to a dampened RNAi effect (Ceron *et al*. 2007; Lehner *et al*. 2006; Wang *et al*. 2005; Wu *et al*., 2012). The reduced HTA suppression in *lin-35* mutants may be in part due to the known pleiotropic effects of loss of *lin-35,* which generally shows slow growth and reduced fertility when compared to the *lin-54* mutant (Chi *et al*. 2006, Kudron *et al*. 2013, Rual *et al*. 2004; J.R.C. and L.N.P data not shown).

The HTA phenotype is a characteristic of not just DREAM complex mutants but also mutants in members of the HP1 complex (Petrella *et al*. 2011). LIN-13 is a member of the HP1 complex, which when mutated shows both ectopic expression of germline genes and the HTA phenotype (Coustham *et al*. 2006; Petrella *et al*. 2011). To determine if the loss of TF HTA suppressors suppresses the HTA phenotype when HP1 complex function is compromised, we performed RNAi against the 15 TF HTA suppressors in a *lin-13* mutant background. RNAi against the 15 TF HTA suppressors failed to significantly suppress the HTA phenotype in *lin-13* mutants (Figure 1c). Together, the combined results of *lin-54*, *lin-35,* and *lin-13* mutants suggest that TF HTA suppressors are acting to drive gene misexpression when the DREAM complex is compromised, but not when the HP1 complex is compromised.

### TF HTA suppressors are required for PGL-1 misexpression in the soma of lin-54 mutants

P-granule proteins, exclusively expressed in the germline of the wild-type, are ectopically expressed in somatic cells of DREAM complex mutants (Wang *et al*. 2005; Petrella *et al*. 2011). To determine if TF HTA suppressors are necessary for the ectopic P-granule expression seen in DREAM complex mutants, we performed RNAi against the 15 TF HTA suppressors in a *lin-54* mutant background and measured the level of the P-granule protein PGL-1 in all of the soma of L1 larvae at 26°C. The normalized mean somatic PGL-1 intensity for each worm was calculated (referred to as adjusted PGL-1 intensity) for a given RNAi treatment (Materials and methods). RNAi of nine of 15 TF HTA suppressors was able to significantly lower the adjusted PGL-1 intensity in *lin-54* mutants at 26°C as compared to L4440 empty vector control (Figure 2 and Table 1). RNAi knock-down of each TF HTA suppressor generally resulted in only a subset of the L1s in the population showing an adjusted PGL-1 intensity in the same range as is seen in wild type worms. For three TF HTA suppressors, *mig-5, ztf-1,* and *dmd-6,* more than half of the L1s showed an adjusted PGL-1 intensity in the wild type range (Figure 2a). The incomplete penetrance seen with suppression of ectopic PGL-1 expression mirrors the partial suppression of the HTA phenotype where a proportion of worms grow to the L4/Adult stage while others are arrested at the L1 stage.

**Figure 2:**
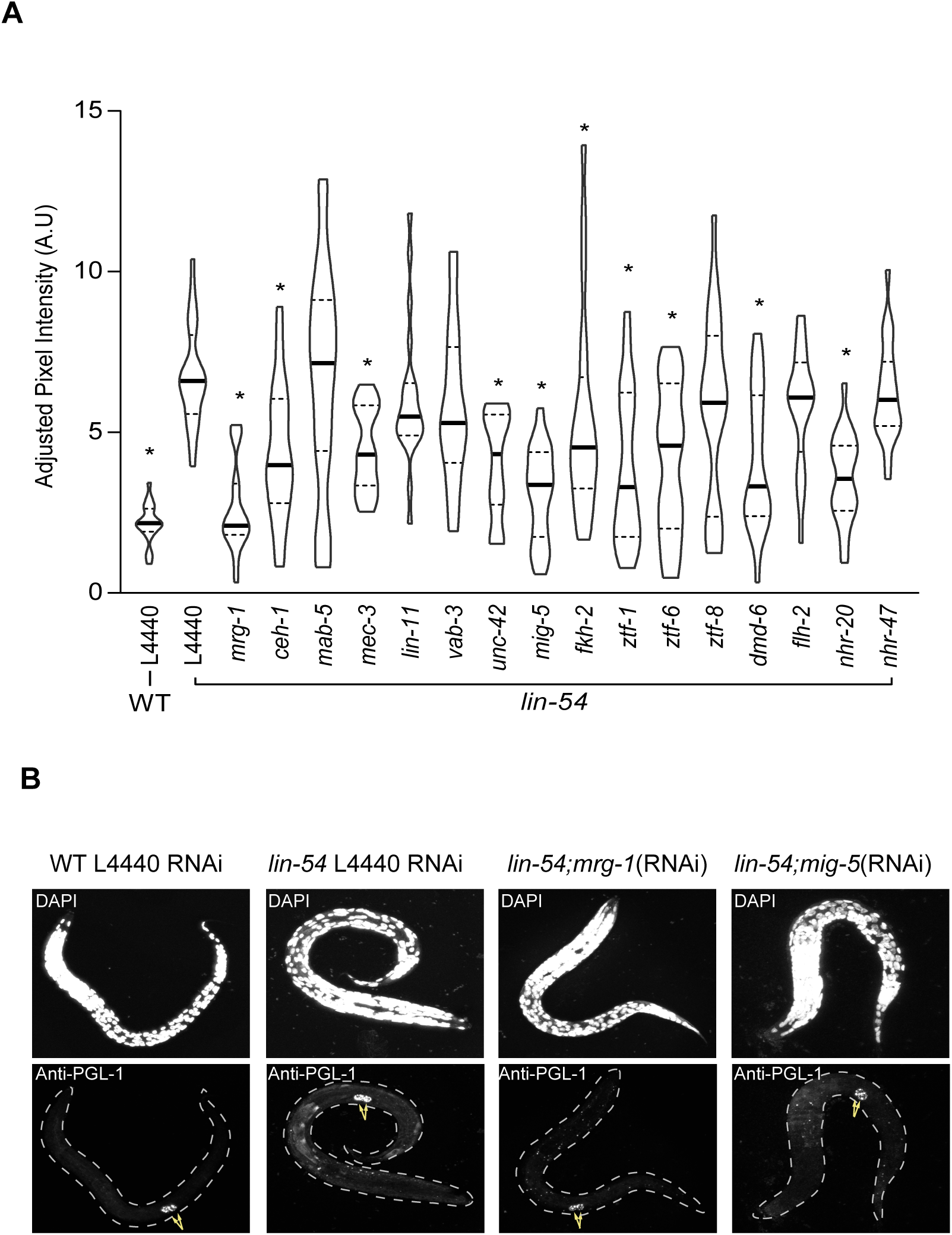
RNAi knock down of 9 transcription factor candidates in a *lin-54* mutant background at 26°C shows reduced ectopic PGL-1 expression. (A) Knock-down of 9 of 15 TF candidates tested showed a significant reduction in adjusted PGL-1 intensity somatic cells of *lin-54* mutants at 26°C when compared to *lin-54* mutants grown on L4440 empty vector bacteria. Y axis indicates the adjusted PGL-1 intensity. The violin plot shows the adjusted PGL-1 intensity frequency distribution for a population of L1s for a particular RNAi treatment. Kolmogorov-Smirnov test was done to compare empty vector L4440 in *lin-54* mutants with every TF. (\**P* value ≤ 0.05). The horizontal solid lines and the dotted lines indicate the median and quartile of the distribution respectively. (B) Representative images for the negative control (L4440 RNAi in a *lin-54* mutant background), positive controls (L4440 RNAi in WT, *mrg-1* RNAi in a *lin-54* mutant background) and a representative sample (*mig-5* RNAi in a *lin-54* mutant background).

In our control experiments, we observed that the adjusted PGL-1 intensity in somatic cells of *lin-54* mutants at 26°C is significantly higher than the level in *lin-54* mutants at 20°C (Figure S2). The adjusted PGL-1 intensity levels in the soma of *lin-54* mutants at 20°C is not discernible from the wild type at either 20°C or 26°C, indicating the limitation of immunostaining in identifying potential subtle differences in adjusted PGL-1 intensity levels between wild type and *lin-54* mutants at 20°C (Figure S2).

### Wnt signaling modulates the HTA phenotype in lin-54 mutants

To determine if the TF HTA suppressors have related functional characteristics, we performed a functional enrichment analysis of the 15 TF candidates (Materials and Methods). Analysis using gProfiler g:GOSt functional profiling revealed overrepresentation of transcription factors involved in the Wnt signaling pathway. Further analysis using PANTHER Overrepresentation Test, resulted in 53 over-represented Gene Ontology (GO) biological process terms. The four most enriched GO terms were neuron fate specification, regulation of animal organ morphogenesis, regionalization, and positive regulation of transcription by RNA polymerase II. Neuronal fate decisions and regionalization are both known to be regulated by Wnt signalling (Bielen *et al*. 2014; Mulligan and Cheyette 2017; Zwamborn *et al*. 2018). Additionally, five of the TF HTA suppressors, including LIN-11, MAB-5, LIN-11, VAB-3, and ZTF-6, have previously been shown to have interaction with Wnt signaling (Doitsidou *et al*. 2018; Gupta and Sternberg 2002; Johnson and Chamberlin 2008, Maloof *et al*. 1999).

We were particularly interested in the possible role of Wnt signaling in DREAM complex mutant phenotypes, as previous data had shown an anterior-to-posterior (A-P) pattern to chromatin compaction defects in DREAM complex mutants (Costello *et al*. 2019). Wnt signaling is important to establish the A-P axis during development and Wnt ligands are found in gradients along the A-P axis (Schroeder *et al*. 1998). To assess if the Wnt signaling pathway plays a role in regulating the HTA phenotype, we performed RNAi against 41 known Wnt signaling associated genes (Sawa *et al*. 2013) and scored for HTA suppression in a *lin-54* mutant background (Figure 3a). RNAi against eight of the Wnt signaling genes resulted in embryonic lethality in both *lin-54* and wildtype animals (Table S1). RNAi against an additional nine Wnt signaling genes resulted in embryonic lethality in only the *lin-54* mutants (Table S1). This may be due to DREAM complex mutants being more sensitive than wild type to perturbations in Wnt signaling or *lin-54* mutants may have enhanced somatic RNAi similar to *lin-35* mutants (Ceron *et al*. 2007; Lehner *et al*. 2006; Wang *et al*. 2005; Wu *et al*., 2012). RNAi against 14 of the 24 remaining Wnt signaling genes significantly suppressed the HTA phenotype in *lin-54* mutants (Figure 3b and Table S1); these are found at multiple levels of Wnt signaling cascade, from Wnt ligands to cytoplasmic factors (Figure 3a). Knockdown of three genes that function in Wnt ligand production and secretion, *mom-1, vps-29,* and *snx-3,* resulted in mild yet significant suppression of the HTA phenotype in *lin-54* mutants (Figure 3b). Knock-down of two embryonically viable Wnt ligands, *egl-20* and *cwn-2,* resulted in suppression of HTA (Figure 3b). EGL-20 is highly expressed in the posterior tail region in wild type animals (Pan *et al*. 2006). Mis-expression of EGL-20 in *lin-54* mutants could be associated with aberrant signaling leading to the HTA phenotype. However, we found that the EGL-20:GFP expression pattern in *lin-54* mutants was limited to the posterior end of the L1 stage worm similar to the wild type pattern at both 20°C and 26°C (Figure S3) (Whangbo *et al*. 1999). Among the transmembrane Wnt receptors tested, only RNAi against *lin-18* suppressed the HTA phenotype to significant levels (Figure 3b). Additionally, RNAi against a number of genes encoding cytoplasmic transducers of the Wnt signal suppressed the HTA phenotype, including two Disheveled orthologs, *mig-5* and *dsh-1*, and one Axin ortholog, *pry-1* (Figure 3b). MOM-4 and TAP-1 are MAP kinase components that function to antagonize Wnt signaling by phosphorylation and inhibition of TCF, a Wnt downstream effector (Smit *et al*. 2004). We observed that knock-down of *tap*-1 suppressed the HTA phenotype. In *lin-54* mutants, knock-down of *mom-4* in resulted in highly penetrant embryonic lethality. Only 2-4 larvae per experiment escaped passed embryonic stage, but all of those grew past L1 arrest stage. The small sample size did not permit significance testing (Figure 3b). Interestingly, RNAi against all three non-canonical Wnt planar cell polarity (PCP) genes, *vang-1*, *fmi-1,* and *prkl-1,* resulted in strong HTA suppression compared to L4440 empty vector RNAi (Figure 3b). Because RNAi against most of the downstream factors controlling Wnt pathway target gene expression were embryonically lethal, we could not assess their role in regulating the HTA phenotype (Table S1). Overall, a specific subset of Wnt signaling genes, found at multiple levels of the signaling cascade, showed suppression of the HTA phenotype.

**Figure 3.**
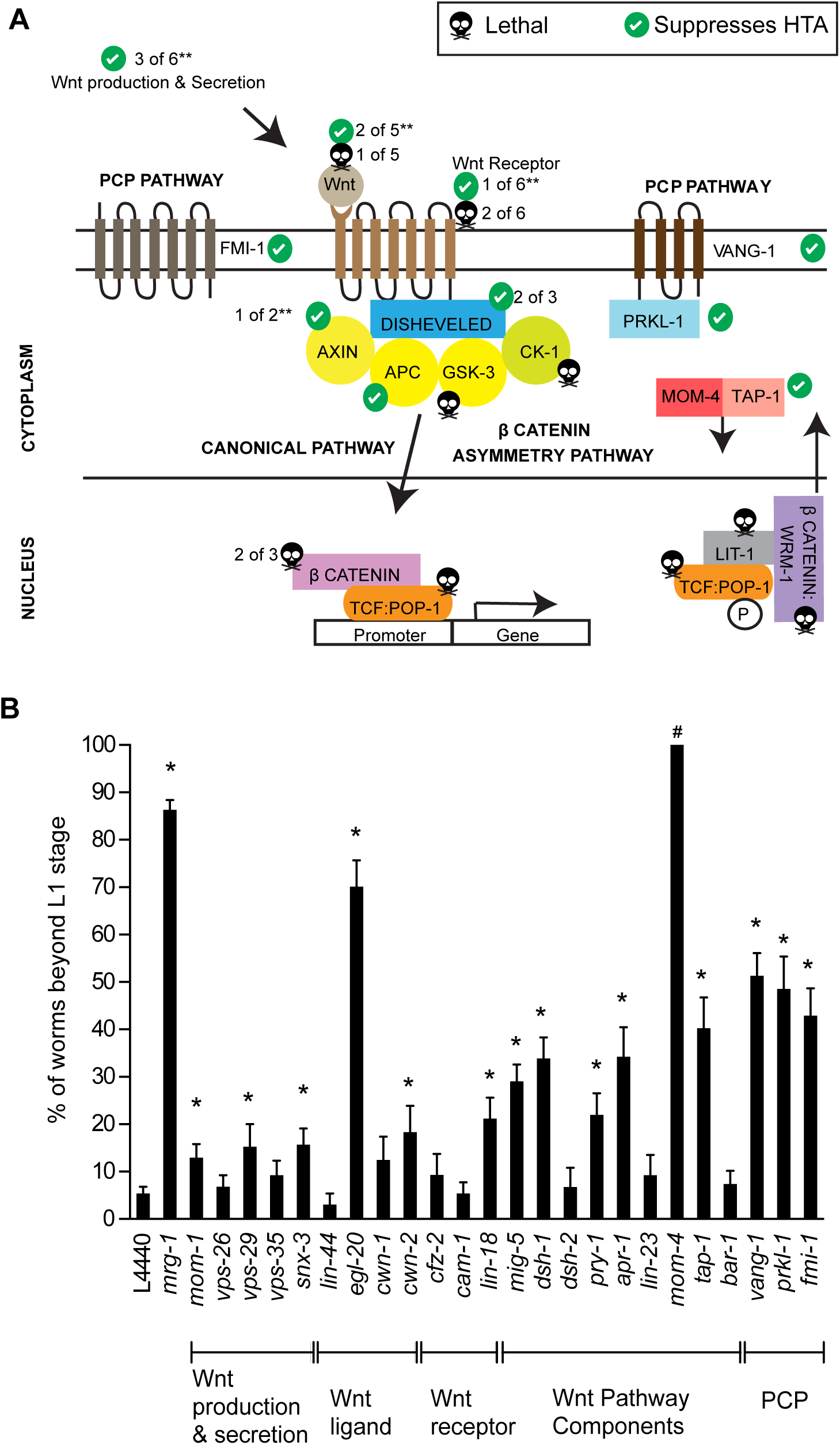
Fourteen Wnt signaling pathway components show HTA suppression phenotype in a *lin-54* mutants. (A) Schematic diagram of the Wnt signaling pathway in *C. elegans*. Check mark on a green filled circle and black skull with cross bone sign indicate genes that show suppression of HTA phenotype in *lin-54* mutants and embryonic lethality when knocked-down respectively. ** indicates that some genes were not tested (refer Table S1 for more details). (B) Knock-down of 14 of 41 Wnt pathway genes tested were able to suppress HTA phenotype in *lin-54* mutant background. Worms that had progeny after the knock-down of Wnt pathway genes in *lin-54* mutants are displayed in the graph. ^#^RNAi of *mom-4* was almost completely embryonic lethal and gave rise to only 2-4 larvae per experiment, but all of those grew past L1 arrest stage. The number of worms scored for *mom-4* was too small a sample size to determine significance. Fisher’s Exact Test was done to compare every Wnt gene RNAi with empty vector L4440. (\**P* value ≤0.05). Error bars indicate standard error of proportion.

### The conserved Wnt/PCP component VANG-1 is required for the HTA phenotype and small brood size of *lin-54* mutants

The strong suppression of the HTA phenotype in *lin-54* mutants with the knock-down of any of the three Wnt PCP pathway components (*vang-1, prkl-1,* and *fmi-1)* led us to further investigate the genetic interaction of the DREAM complex and Wnt/PCP pathways. VANG-1 is the highly conserved C. elegans homolog of PCP pathway associated transmembrane Vangl protein necessary for mediating PCP signaling (He *et al*. 2018; Honnen *et al*. 2012; Yang & Mlodzik 2015). We crossed the *vang-1*(ok1142) mutation, which lacks 162 amino acids of the C-terminus (Honnen *et al*. 2012), to the *lin-54* mutant to create a *vang-1; lin-54* double mutant. We found that the double mutant showed a complete suppression of the HTA phenotype (Figure 4a). Additionally, the adjusted PGL-1 intensity distribution in somatic cells of lin*-54; vang-1* mutants was significantly reduced in comparison to *lin-54* mutants at 26°C (Figure 4b).

**Figure 4.**
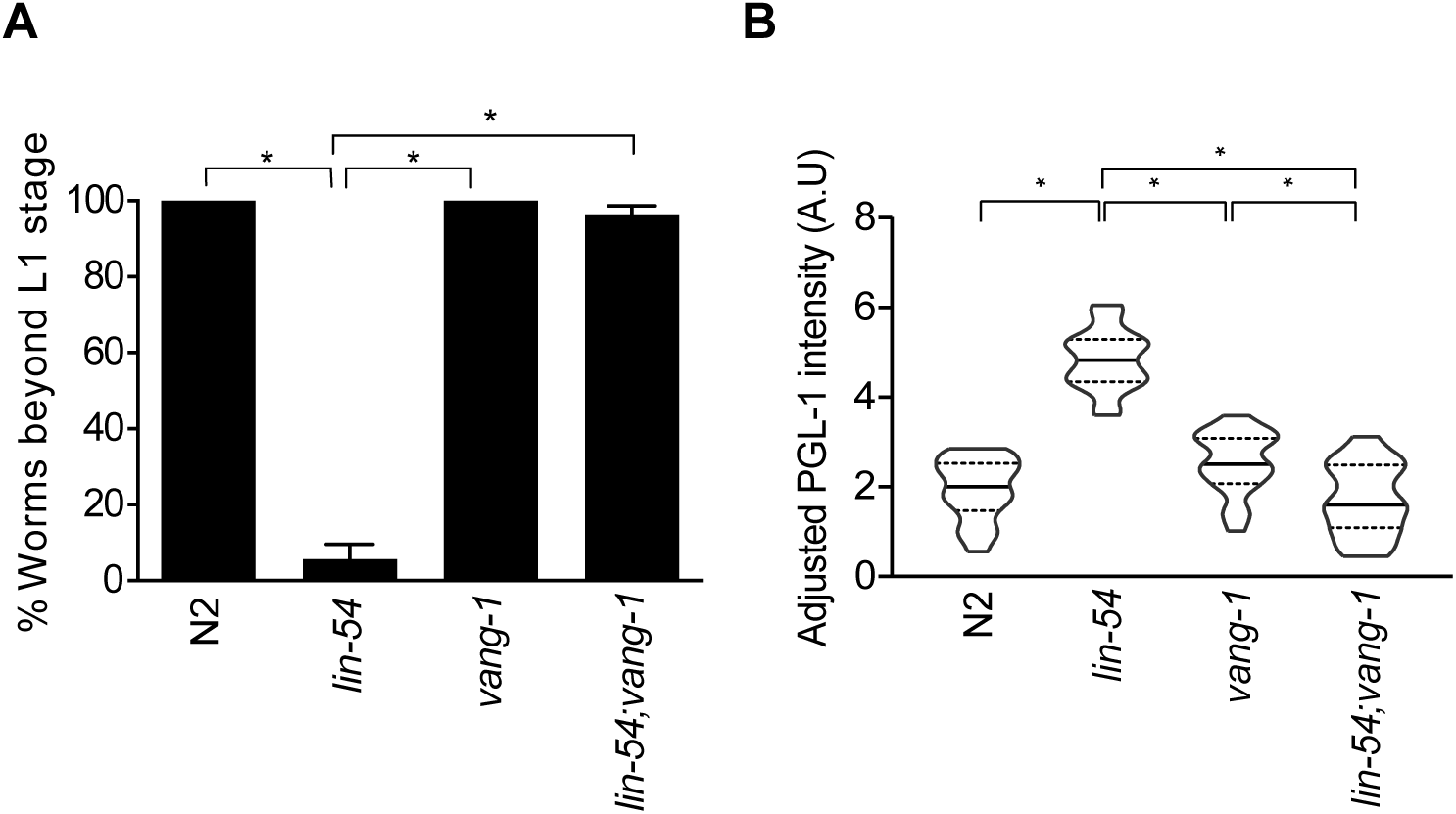
*lin-54; vang-1* mutants completely suppresses the HTA phenotype and shows reduced ectopic PGL-1 expression close to wild-type levels. (A) Wild type, *vang-1* and *lin-54; vang-1* mutants do not show the HTA phenotype and grow to become adults in contrast to *lin-54* mutants that arrested at the L1 stage at 26°C. (Fisher’s Exact test \**P* value ≤ 0.05). Error bars indicate standard error of proportion. (B) *lin-54; vang-1* mutants show reduced somatic adjusted PGL-1 intensity at 26°C comparable to wild-type levels. Y-axis indicates adjusted PGL-1 intensity. The violin plot shows the adjusted PGL-1 intensity frequency distribution for a population of L1s under a particular RNAi treatment. The horizontal solid lines and dotted lines indicate the median and quartile of the distribution respectively. (Kolmogorov-Smirnov test \**P* value ≤ 0.05).

When analyzing the suppression of HTA, we observed that *lin-54; vang-1* double mutant worms at 26°C were fertile. Mutations in most previously identified HTA suppressors, such as *mrg-1* and *mes-4,* result in sterility, (Fujita *et al*. 2002; Garvin *et al*. 1998, Petrella *et al*. 2011). We analyzed the brood size of hermaphrodites under three different temperature regimes: hermaphrodites kept at 20°C continuously, hermaphrodites raised at 20°C until the L4 stage that were then up-shifted to 26°C, and hermaphrodites kept at 26°C continuously. *lin-54* single mutants have significantly smaller mean brood sizes than wild type or *vang-1* single mutants at both 20°C and when up-shifted to 26°C (Figure 5). Due to the HTA phenotype, *lin-54* single mutants could not be analyzed when raised at 26°C. The smaller brood size seen in *lin-54* single mutants is completely suppressed in *lin-54; vang-1* double mutants. In fact, double mutants showed significantly larger mean brood sizes than wildtype hermaphrodites at all three temperatures tested (Figure 5). In summary, loss of *vang-1* produced almost complete suppression of the *lin-54* HTA phenotype, ectopic germline gene expression, and fertility defects.

**Figure 5.**
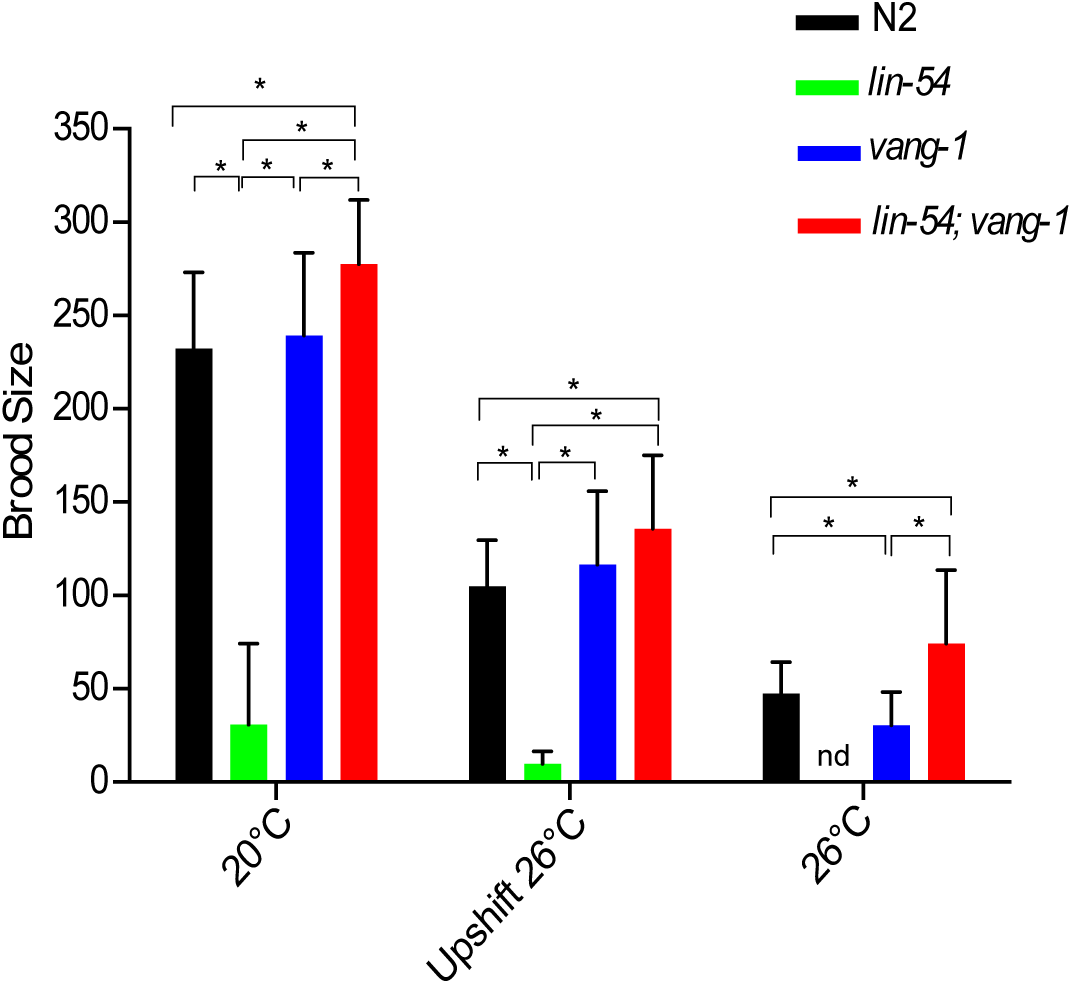
*lin-54;vang-1* mutants suppresses small brood size phenotype of *lin-54* mutants. The *lin-54; vang-1* mutant showed a significantly larger brood size compared to wild type, *lin-54,* and *vang-1* mutants at 20°C and 26°C. (Welch’s T test \**P* value ≤ 0.05). Error bars indicate standard deviation.

## DISCUSSION

Loss of the DREAM complex results in misexpression of germline genes in somatic cells and high temperature larval arrest (Petrella *et al*. 2011; Wang *et al*. 2005; Wu *et al*. 2012). Using an RNAi suppressor screen, we identified a set of 15 TFs that contribute to the HTA phenotype. Additionally, we found that genes that function in Wnt signaling, notably those in the Wnt/PCP pathway, are also required for the high temperature larval arrest. Based on these results, we propose a model whereby embryonic TFs and the Wnt/PCP pathway each contribute to ectopic activation of germline genes, and the resulting larval arrest, in DREAM complex mutants.

The HTA phenotype of DREAM complex mutants has been attributed to significant germline gene misexpression in embryonic intestinal cells, leading to intestinal cell dysfunction and nutritional deprivation even in the presence of food (Petrella *et al*. 2011). There are two potential sources for the TFs that we found contribute to germline gene expression in DREAM complex mutants. First, these TFs may be normally expressed in wild-type embryonic somatic tissues, but unable to bind germline gene targets in the presence of the DREAM complex. For this class of TFs, activation of germline genes may result from their recruitment to novel gene targets in the absence of the DREAM complex. Second, the TFs themselves could be genes normally repressed by the DREAM complex, and are ectopically expressed in DREAM complex mutants. Analysis of single-cell RNA-seq data from the *C. elegans* embryo showed that all but one of the TF HTA suppressors found in our screen are normally expressed in the wild-type embryonic intestinal lineage (Packer *et al*. 2019), a finding consistent with the former possibility. The sole exception is *ceh-1,* which is expressed embryonically only in the anal sphincter muscles, AVK neurons, and anterior body wall muscles but with no clear evidence of intestinal expression (Packer *et al*. 2019). Taken together, these data suggest that in the absence of the DREAM repressor complex, TFs normally expressed in embryonic somatic cells can be coopted to bind new genomic locations, leading to gene misexpression and cell fate changes.

In our RNAi screen, we tested all TFs that were capable of binding to DREAM complex target genes in a Y1 Hybrid assay (Reece-Hoyes *et al*. 2011). While these proteins are predicted to function as transcription factors based on the presence known DNA binding domains, they have not necessarily been validated as TFs *in vivo.* In the Y1 hybrid data set, MIG-5 bound 50% of the DREAM target genes that were tested, the largest number of all the TF HTA suppressors (Table 1). Knockdown of *mig-5* showed strong suppression of the HTA phenotype and ectopic PGL-1 expression. However, MIG-5 is not a known transcription factor, but instead one of three worm Disheveled (Dsh) proteins, which all contain a winged-helix DNA binding domain. Dsh proteins function in Wnt signaling by stabilizing activated β-catenin in the cytoplasm (Figure 3a) (Jackson *et al*. 2012; Salic *et al*. 2000). Dsh proteins are also known to be an important branch point between the canonical Wnt and Wnt/PCP pathways (Yan *et al*. 2001; Salic *et al*. 2000; Walllingford *et al*. 2005). In some organisms Dsh proteins shuttle to the nucleus, which is has been shown to be necessary for Wnt signaling (Cheyette *et al*. 2002; Habas *et al*. 2005; Itoh *et al*. 2005; Torres *et al*. 2000). Furthermore, in mammals Dsh proteins have been shown to interact directly with transcription factors within the nucleus (Gan *et al*. 2008; Wang *et al*. 2015). Nuclear localization of MIG-5 has not been observed in *C. elegans;* however, the known nuclear localization signal of *Xenapus* Dsh is almost completely conserved within MIG-5 (Itoh *et al*. 2005). Further analysis is needed to determine if we have uncovered a novel role for MIG-5 in the nucleus in *C. elegans* or if MIG-5 is solely acting by transducing non-canonical Wnt/PCP signaling in DREAM complex mutants.

One of the striking findings from our screens is the substantial role that Wnt/PCP signaling plays in driving the HTA and germline gene expression phenotypes in *lin-54* mutants. Many components of the Wnt signaling cascade can differ between tissues and developmental times, which in turn can lead to different outcomes depending on the specific cellular context (Eisenmann 2005; Hardin *et al*. 2008; Kwon *et al*. 2008; Sokol 2015). Therefore, it is unclear if the suppression of the HTA phenotype seen with loss of other Wnt signaling components is due to the specific loss of canonical Wnt signaling or Wnt/PCP signaling. However, our data supports an important role for Wnt/PCP signaling in the HTA phenotype. First, loss of any of the three components specific to the Wnt/PCP pathway, *vang-1, prkl-1,* and *fmi-1,* all lead to significant suppression of HTA. Second, some of the other Wnt signaling components that suppress the HTA phenotype have been shown to interact with the Wnt/PCP pathway. Most strikingly is the strong suppression seen with knockdown of the Wnt ligand *egl-20*. Of the five Wnt ligands, EGL-20 has most often been shown to interact with the Wnt/PCP pathway (Green *et al*. 2008; Mentink *et al*. 2018). In the intestine, the only documented role of Wnt/PCP signaling is for proper orientation of intestinal cells during intestinal morphogenesis (Asan *et al*. 2016; Hoffmann *et al*. 2010), but in other tissues Wnt/PCP signaling can help to regulate A-P fate specification (Ackley 2014; Antic *et al*. 2010; He *et al*. 2018; Mattes et al. 2018; Mentink *et al*. 2018). It is still unclear what role Wnt/PCP signaling plays in regulating gene expression and fate specification in the intestine.

How could the Wnt/PCP pathway be driving the HTA phenotype? We propose two non-mutually exclusive models of how Wnt/PCP signaling could drive ectopic DREAM complex target expression leading to HTA. The first model is that Wnt/PCP signaling may be necessary for the expression of the transcription factors that directly bind to DREAM target loci and drive their ectopic expression. The proper asymmetric anterior-posterior (A-P) expression of three of the TF HTA suppressors found in our screen, MAB-5, LIN-11, and VAB-3, requires Wnt signaling in neuronal and vulval tissues (Gupta and Sternberg 2002; Johnson and Chamberlin 2008, Maloof *et al*. 1999). A fourth TF HTA suppressor, MEC-3, also has known asymmetric A-P expression, although this pattern has yet to be attributed specifically to Wnt signaling (Way *et al*. 1992). The second model is that Wnt/PCP signaling regulates the A-P patterning of the intestine, and that this pattern is itself a necessary state for ectopic expression of DREAM target loci. While chromatin structure in wild-type embryonic cells undergoes rapid compaction, we previously showed that this process is delayed in DREAM complex mutants, and shows an A-P pattern where anterior cells are more severely delayed (Costello and Petrella 2019). A-P axis patterning in *C. elegans* is primarily established by Wnt signaling (Lin *et al*. 1998; Schroeder *et al*. 1998). Therefore, underlying aspects of a A-P pattern established by Wnt signaling may facilitate the delayed chromatin compaction in DREAM complex mutants at 26°C, which in turn allows TFs to bind and activate germline genes. Distinguishing between these two models will require further analysis of the interaction of Wnt signaling, chromatin compaction, and TF expression in DREAM complex mutants.

The DREAM complex, ectopic expression of germline genes, and Wnt/PCP signaling all have known roles in cancer acquisition and progression. The DREAM complex is completely conserved in mammals, and loss of DREAM complex function is associated with cancers (Engeland 2018; Patel *et al*. 2019; Reichert *et al*., 2010). Some tumors are known to ectopically express a group of germline specific genes called cancer-testis (CT) genes. CT gene expression has been shown to correlate with increased proliferative index, higher tumor grade, and poorer response to therapies in many studies and across a broad range of tumor types (Gure *et al*., 2005; Wang *et al*., 2016; Whitehurst, 2014; Xu *et al*., 2014; Yakirevich *et al*., 2003). Wnt/PCP signaling is important in cancer progression and metastasis in a number of tumor types including lung cancer, melanoma, and breast cancer (Humphreis and Mlodzik 2018; Katoh 2005). Our data indicate that these three pathways are linked in *C. elegans* where Wnt signaling via the non-canonical PCP mechanism leads to activation of germline genes in the absence of DREAM complex binding. The identification of the upstream VANG-1 of Wnt/PCP pathway provides a useful starting point to help dissect out the intermediate signaling molecules to activate germline genes in somatic cells of DREAM complex mutants. Our findings, provide insight into potential pathways that are important in the activation of germline genes in cancer cells that could be targets to limit cancer progression.

## ACKNOWLEDGEMENTS

We would like to thank our undergraduate student Katherine Adams for helping with RNAi screening of some transcription factor candidates. Thanks to CGC, which is funded by NIH Office of Research Infrastructure Programs (P40 OD010440), for providing some of the strains for experiments. We appreciate the time and efforts taken by Dr. Anita Manogaran & Dr. Claire de la Cova for comments on the manuscript.

---

**Figure S1:**
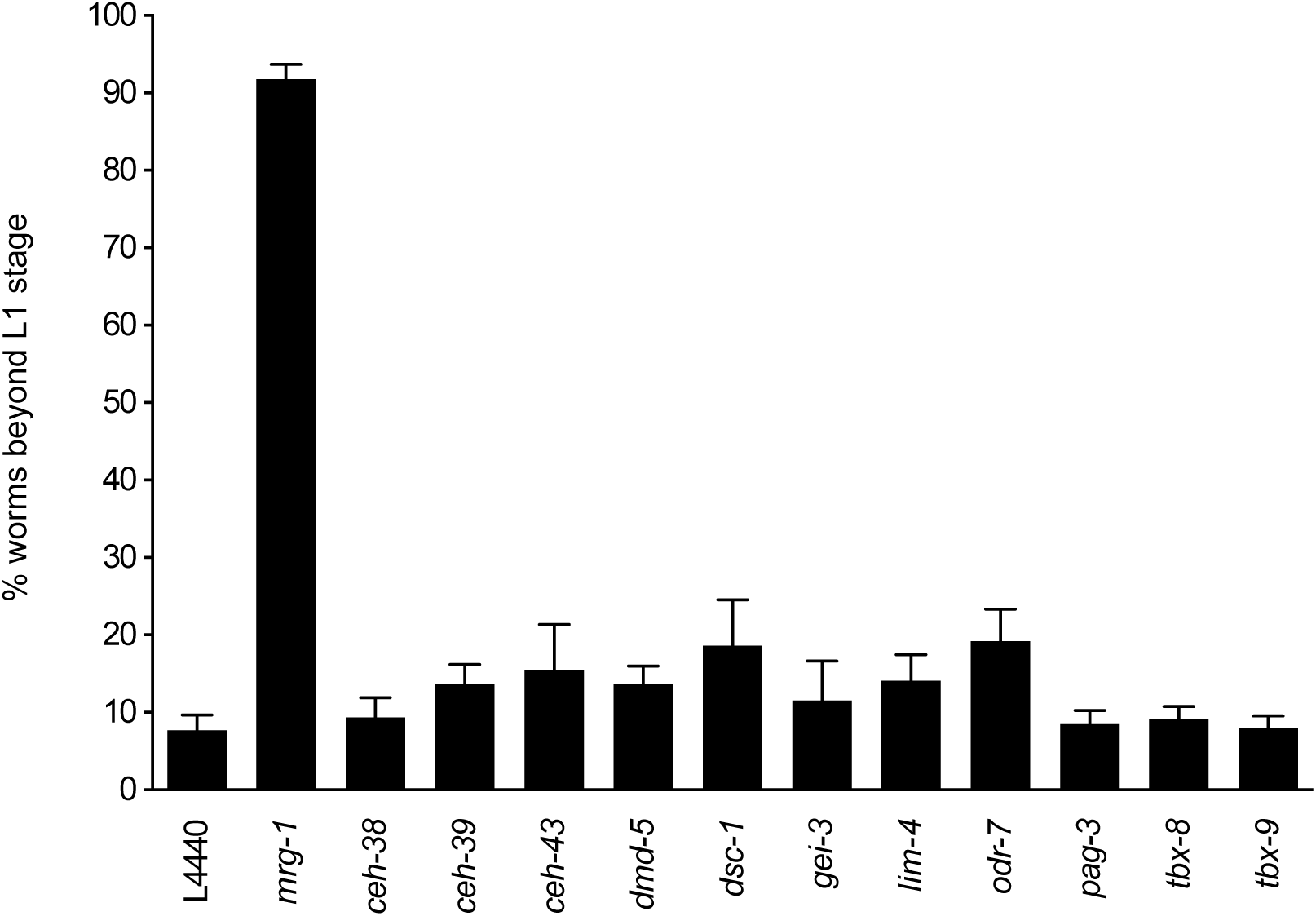
Eleven Transcription Factors show low levels of HTA phenotype suppression. Knock-down of 11 of 26 transcription factor candidates tested showed suppression of the HTA phenotype in *lin-54* mutants but did not show a significant difference from L4440 empty vector RNAi. Error bars indicate standard error of proportion.

**Figure S2:**
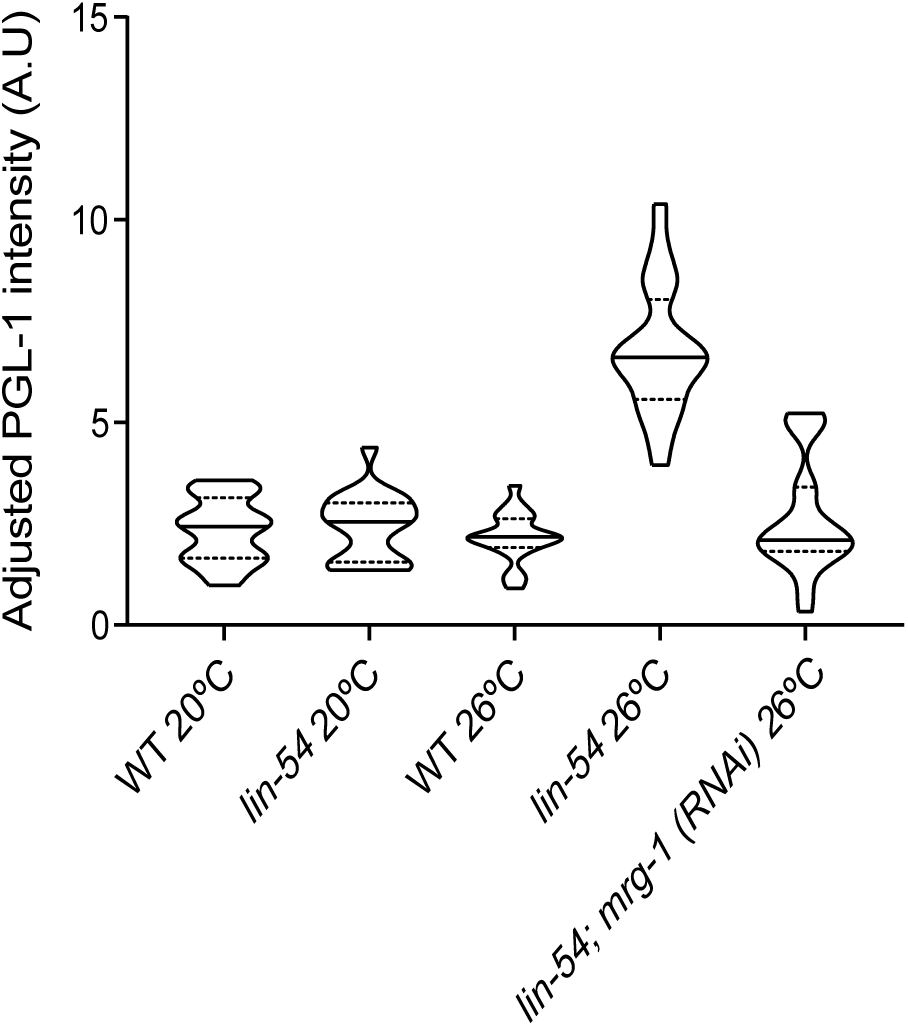
Frequency distribution of somatic adjusted PGL-1 intensity of control samples. Wild type and *lin-54* mutants were grown on L4440 empty vector RNAi bacteria unless otherwise indicated. Y-axis indicates adjusted PGL-1 intensity. The violin plot shows the adjusted PGL-1 intensity frequency distribution for a population of L1s under a particular RNAi treatment. The horizontal solid lines and dotted lines indicate the median and quartile of the distribution respectively. Wild type at 20°C, wild type at 26°C and *lin-54* mutants at 20°C do not show a significant different level of ectopic PGL-1 expression from each other but all the three showed a significant difference in ectopic PGL-1 expression from *lin-54* mutants at 26°C. Kolmogorov-Smirnov test was used for testing significance and the *P value cut off is 0.05.

**Figure S3:**
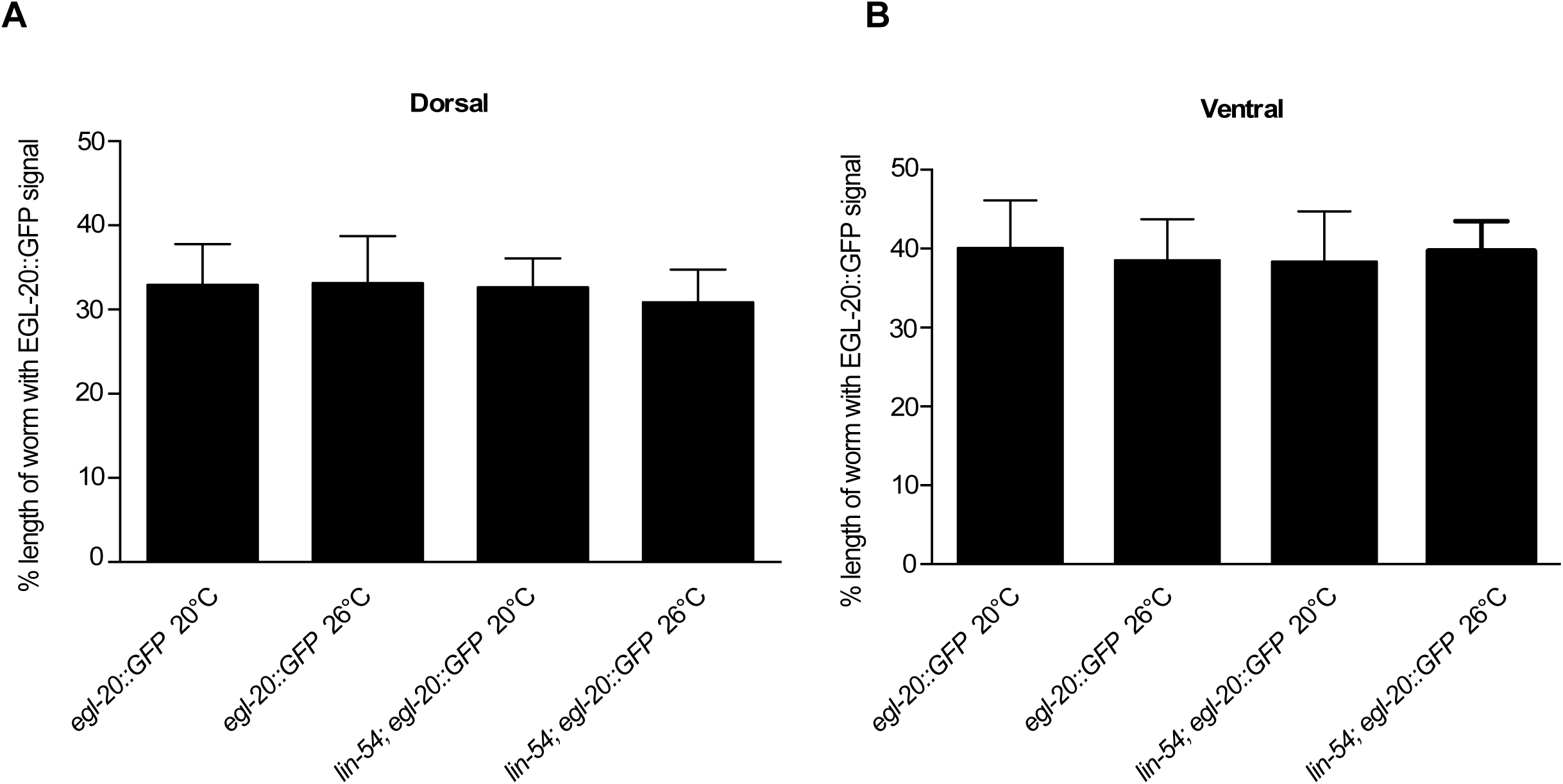
Wnt ligand EGL-20 localization is not altered in *lin-54* mutants. The Wnt EGL-20 gradient was measured by determining the linear distance of GFP on both dorsal (left panel) and ventral side (right panel) from the posterior end of the worm divided by total length of the worm. No significant difference was seen between wild type and *lin-54* mutants at either 20°C or 26°C.

**Table S1:** Results of RNAi knock-down of Wnt signaling components.

| Gene Name | Function# | %HTA | SEM | RNAi lethality in WT | RNAi lethality in *lin-54* mutants |
| --- | --- | --- | --- | --- | --- |
| *mom-1* | Wnt production & secretion | 12.98 | 2.84 | No | No |
| *vps-26* | Wnt production & secretion | 6.85 | 2.42 | No | No |
| *vps-29* | Wnt production & secretion | 15.28 | 4.72 | No | No |
| *vps-35* | Wnt production & secretion | 9.27 | 3.06 | No | No |
| *snx-3* | Wnt production & secretion | 15.69 | 3.44 | No | No |
| *mig-14* | Wnt production & secretion | NA | NA | ND* | ND* |
| *mom-2* | Wnt ligand | NA | NA | No | Yes |
| *lin-44* | Wnt ligand | 3.04 | 2.34 | No | No |
| *egl-20* | Wnt ligand | 70.13 | 5.55 | No | No |
| *cwn-1* | Wnt ligand | 12.44 | 4.98 | No | No |
| *cwn-2* | Wnt ligand | 18.33 | 5.58 | No | No |
| *sfrp-1* | Wnt Inhibitor | NA | NA | ND* | ND* |
| *mom-5* | Wnt receptors | NA | NA | No | Yes |
| *cfz-2* | Wnt receptors | 9.33 | 4.39 | No | No |
| *cam-1* | Wnt receptors | 5.37 | 2.39 | No | No |
| *lin-17* | Wnt receptors | NA | NA | Yes | Yes |
| *lin-18* | Wnt receptors | 21.18 | 4.43 | No | No |
| *mig-1* | Wnt receptors | NA | NA | ND* | ND* |
| *mig-5* | Dishevelled | 29.05 | 3.57 | No | No |
| *dsh-1* | Dishevelled | 33.88 | 4.41 | No | No |
| *dsh-2* | Dishevelled | 6.73 | 4.07 | No | No |
| *pry-1* | $\beta$ Catenin Destruction Complex | 21.93 | 4.6 | No | No |
| *apr-1* | $\beta$ Catenin Destruction Complex | 34.27 | 6.18 | No | No |
| *lin-23* | $\beta$ Catenin Destruction Complex | 9.26 | 4.27 | No | No |
| *gsk-3* | $\beta$ Catenin Destruction Complex | NA | NA | Yes | Yes |
| *kin-19* | $\beta$ Catenin Destruction Complex | NA | NA | Yes | Yes |
| *axl-1* | $\beta$ Catenin Destruction Complex | NA | NA | ND* | ND* |
| *mom-4* | Tak | 100 | 0 | No | No |
| *tap-1* | Tab | 40.25 | 6.5 | No | No |
| *lit-1* | Nlk | NA | NA | No | Yes |
| *bar-1* | $\beta$ Catenin | 7.37 | 2.79 | No | No |
| *hmp-2* | $\beta$ Catenin | NA | NA | No | Yes |
| *sys-1* | $\beta$ Catenin | NA | NA | Yes | Yes |
| *wrm-1* | $\beta$ Catenin | NA | NA | Yes | Yes |
| *unc-37* | Groucho | NA | NA | Yes | Yes |
| *pop-1* | TCF | NA | NA | Yes | Yes |
| *vang-1* | Wnt/PCP member | 51.3 | 4.79 | No | No |
| *prkl-1* | Wnt/PCP member | 48.53 | 6.87 | No | No |
| *fmi-1* | Wnt/PCP member | 42.9 | 5.75 | No | No |
| *hda-1* | Wnt signaling interactors | NA | NA | No | Yes |
| *skn-1* | Wnt signaling interactors | NA | NA | No | Yes |
| *med-1* | Wnt signaling interactors | NA | NA | No | Yes |
| *rho-1* | Wnt signaling interactors | NA | NA | No | Yes |
| *let-502* | Wnt signaling interactors | NA | NA | No | Yes |
| *pal-1* | Wnt signaling interactors | NA | NA | Yes | Yes |

